# Highly efficient *Runx1* enhancer eR1-mediated genetic engineering for fetal, child and adult hematopoietic stem cells

**DOI:** 10.1101/2021.11.24.469958

**Authors:** Cai Ping Koh, Avinash Govind Bahirvani, Chelsia Qiuxia Wang, Tomomasa Yokomizo, Cherry Ee Lin Ng, Linsen Du, Vinay Tergaonkar, Dominic Chih-Cheng Voon, Hiroki Hosoi, Takashi Sonoki, Mok Meng Huang Michelle, Akiko Niibori-Nambu, Yi Zhang, Archibald S. Perkins, Zakir Hossain, Daniel G. Tenen, Yoshiaki Ito, Byrappa Venkatesh, Motomi Osato

**Author notes:** Corresponding Authors: Motomi Osato, MD, PhD, Cancer Science Institute of Singapore, National University of Singapore, 14 Medical Drive, Singapore 117599.; Byrappa Venkatesh. These authors contributed equally to this work (Cai Ping Koh, Avinash Govind Bahirvani).

## Abstract

A *cis*-regulatory genetic element which targets gene expression to stem cells, termed stem cell enhancer, serves as a molecular handle for stem cell-specific genetic engineering. Here we show the generation and characterization of a tamoxifen-inducible CreER^T2^ transgenic (Tg) mouse employing previously identified hematopoietic stem cell (HSC) enhancer for *Runx1*, eR1 (+24m). Kinetic analysis of labeled cells after tamoxifen injection and transplantation assays revealed that eR1-driven CreER^T2^ activity marks dormant adult HSCs which slowly but steadily contribute to unperturbed hematopoiesis. Fetal and child HSCs which are uniformly or intermediately active were also efficiently targeted. Notably, a gene ablation at distinct developmental stages, enabled by this system, resulted in different phenotypes. Similarly, an oncogenic Kras induction at distinct ages caused different spectrums of malignant diseases. These results demonstrate that the eR1-CreER^T2^ Tg mouse serves as a powerful resource for the analyses of both normal and malignant HSCs at all developmental stages.

## Introduction

There has been considerable interest in identifying a *cis*-regulatory element that targets gene expression to hematopoietic stem cells (HSCs). Characterization of such an enhancer, termed HSC enhancer, holds promise of generating important insights into the transcriptional programs responsible for “stemness”, and will also provide a powerful tool for manipulation of HSCs in experimental animal models, such as mouse and zebrafish. A HSC enhancer, in combination with reporters such as EGFP or LacZ, can be used for the marking of HSCs [1-3]. This way of HSC identification is called genetic marking of HSCs. In addition, the use of fluorescence reporters, like EGFP, enables *in vitro* and *in vivo* live imaging of HSCs. Such a HSC enhancer can also drive the expression of any gene of interest, namely transgenesis in HSCs. Furthermore, when engineered to drive the expression of Cre recombinase gene, a HSC enhancer enables excision of a floxed allele, thereby allowing for genetic engineering, including gene targeting in a HSC-specific manner.

A series of candidate HSC enhancers have been identified. However, none of them have shown activities restricted to HSC compartment, but much broader expression including less immature progenitor fractions and endothelial cells which are major components in the bone marrow (BM) [4-8]. We and others have succeeded in identifying an intronic <u>e</u>nhancer for *<u>R</u>unx<u>1</u>*, named eR1 (+24m), which is highly active in HSCs [2, 9, 10]. To exploit this HSC enhancer activity of eR1, eR1-CreER^T2^ transgenic (Tg) mouse was generated.

## Results

### Generation and selection of eR1-CreER^T2^ Tg mouse lines

eR1 is a regulatory element, being active in HSCs and located at 24 kb downstream of mouse *Runx1* P1 promoter (Figure 1A). A Tg mouse with CreER^T2^ recombinase expression driven by eR1 was generated (Figure 1B). Crossing with a reporter strain, Rosa26-LSL-tdTomato mice, selection of appropriate Tg line was conducted amongst 5 Tg lines, in all of which successful germline transmission of Tg was confirmed (Figure S1A). Line #5-2 was selected for subsequent analyses for normal HSCs due to its highest labeling efficiency, 35.83% Tomato^+^ cells in peripheral blood (PB) at 1 week after a single tamoxifen injection. For malignant HSC studies, another Tg line, #3-17 was selected for its lowest leakage (0.01%) in the absence of tamoxifen treatment (Figure S1A and S1D). Optimization study for tamoxifen dosage and mouse age, using the eR1-CreER^T2^ Tg mouse line #5-2, demonstrated that the highest labeling efficiency with minimal toxicity can be achieved at 0.2 mg/g x 6 times on alternative days from 3 to 4 weeks old Tg mice (Figure 1C-H, S1E and S1F). Using this dosing regimen, no apparent changes were found in hematological parameters including flow cytometry analyses for stem/progenitor fractions.

**Figure 1.**
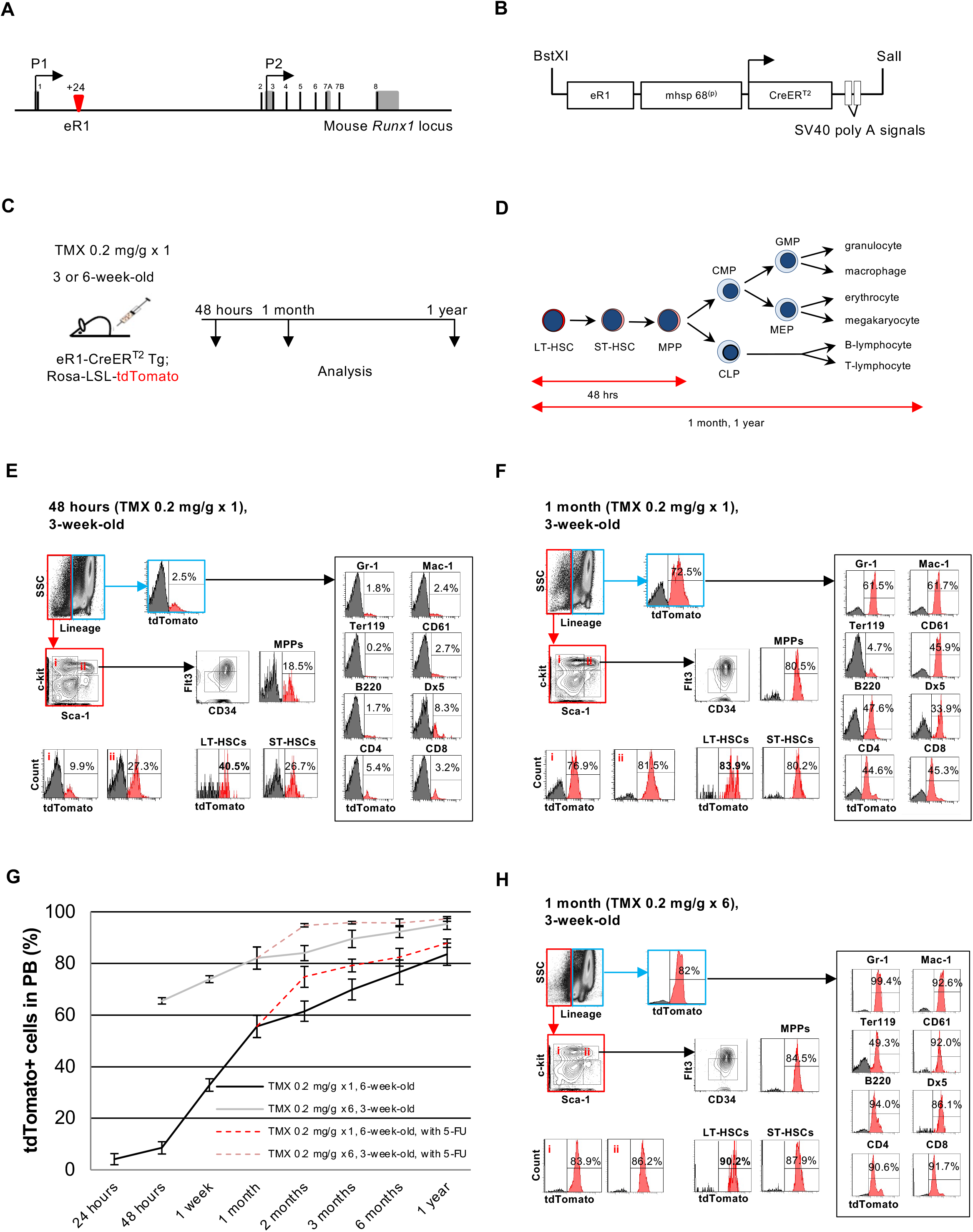
Highly efficient HSC marking in adult and child eR1-CreER^T2^ mice. (A) Schematic diagram of the mouse *Runx1* locus showing eR1 location (red arrow). Black bars represent exons while grey boxes indicate UTR. (B) Transgene construct used to generate the eR1-CreER^T2^ Tg mouse. (C) Schematic diagram demonstrates analysis schedule. 3-week-old or 6-week-old eR1-CreER^T2^ Tg; Rosa26-LSL-tdTomato mice were analyzed at 48 hours (initial target cells), 1 month (lineage tracing) and 1 year (long-term label retaining cells) after TMX injection. (D) Expected dynamics of tdTomato^+^ cells in eR1-CreER^T2^ Tg; Rosa26-LSL-tdTomato mice. (E) Flow cytometry analysis of BM cells from eR1-CreER^T2^ Tg; Rosa26-LSL-tdTomato mouse at 48 hours after TMX injection. Note the highest percentage of tdTomato^+^ cells in CD34^-^Flt3^-^ KSL fraction (LT-HSC). (F) Flow cytometry analysis of BM cells from eR1-CreER^T2^ Tg; Rosa26-LSL-tdTomato mouse at 1 month after TMX injection. All lineage marker –positive cells exhibit tdTomato signals. (G) Long-term kinetics of tdTomato^+^ cells in PB of eR1-CreER^T2^ Tg; Rosa26-LSL-tdTomato mice with single 0.2 mg/g TMX injection into 6-week-old mice (black line) or six times 0.2 mg/g TMX into 3-week-old mice (grey line). 5-FU challenge was conducted at one month after TMX injection (dotted lines). (H) Labeling efficiency in BM at 1 month after TMX injection with indicated dosages to 3-week-old mice. Abbreviations: TMX, tamoxifen; HSC, hematopoietic stem cell; HPC, hematopoietic cell; BM, bone marrow; PB, peripheral blood; 5-FU, 5-fluorouracil; UTR, untranslated region; LT-HSC, long-term hematopoietic stem cell; ST-HSC, short-term hematopoietic stem cell; MPP, multipotent progenitor; CMP, common myeloid progenitor; CLP, common lymphoid progenitor; GMP, granulocyte and macrophage progenitor; MEP, megakaryocyte and erythroid progenitor.

### Highly efficient HSC marking in adult and child eR1-CreER^T2^ Tg mice

In tamoxifen-induced labeling experiment, successful HSC marking was evaluated according to the following three criteria: 1) Initially targeted cells are HSCs, 2) Labeling is traced in all hematopoietic lineages (lineage tracing), and 3) Labeling is maintained for a significantly long period as long-term label-retaining cells. To interrogate these points, analysis of eR1-CreER^T2^ Tg; Rosa26-LSL-tdTomato mice was conducted at 48 hours, 1 month and 1 year after 0.2 mg/g single tamoxifen injection (Figure 1C). It is postulated that if initially targeted cells are HSCs, all the progeny cells derived from HSCs will retain the tdTomato signals in the Rosa26-LSL-tdTomato mouse system. Hence, one month after tamoxifen injection, all hematopoietic cells (HPCs) are expected to display tdTomato signals. The schematic diagram in Figure 1D depicts the expected dynamics of tdTomato^+^ cells in the eR1-CreER^T2^; Rosa26-LSL-tdTomato mice.

In line with the reported eR1 activity in HSCs, the highest marking (40.5% tdTomato^+^ cells) was observed in long term hematopoietic stem cells (LT-HSC) fraction at 48 hours after tamoxifen injection (Figure 1E). The percentages of tdTomato^+^ cells significantly decreased with differentiation in a stepwise manner and only 2.5% lineage positive (Lin^+^) cells expressed tdTomato. These results suggest that the initially targeted cells by eR1-mediated labeling are largely LT-HSCs and, to a lesser extent, short term hematopoietic stem cells (ST-HSCs) and multipotent progenitor (MPP) cells.

At one month after tamoxifen injection, 72.5% Lin^+^ cells demonstrated tdTomato signals (Figure 1F) and labeled cells were observed throughout all hematopoietic lineages, suggesting that originally targeted cell population via eR1 activity were primarily HSCs. The percentage of tdTomato^+^ cells in PB progressively increased until the longest time point of observation, one and half year after tamoxifen injection (Figure 1G).

It has been recently reported that actively proliferating ST-HSCs and MPPs mainly contribute to steady state adult hematopoiesis, whereas dormant LT-HSCs usually do not but serve as a reservoir population for stressed hematopoiesis [11, 12]. To further prove that the dormant adult HSCs were targeted by eR1-CreER^T2^, 5-fluorouracil (5-FU) was injected into eR1-CreER^T2^ Tg; Rosa26-LSL-tdTomato mice one month after tamoxifen injection. In two different injection cohorts, the percentages of tdTomato^+^ cells in PB increased rapidly after 5-FU challenge (dotted lines in Figure 1G). These results suggest that eR1-CreER^T2^ mediated labeled cells include dormant adult LT-HSCs, which minimally contribute to unperturbed adult hematopoiesis but play an active role in stressed hematopoiesis.

It is well recognized that HSC behavior is different at distinct developmental stages. HSC population rapidly expands from E10.5 onwards until 4 weeks after birth, while HSCs are largely maintained in quiescence during the adult stage [13]. LT-HSCs were highly proliferative in 3-week-old mice as the percentage of tdTomato^+^ cells in LT-HSCs increased from 40% to 70% in the period from 48 hours to 1 week after tamoxifen injection, whereas the percentage of tdTomato^+^ cells increased marginally (45% to 50%) in LT-HSCs from 6-week-old mice, concordant with the notion that HSCs are maintained in a quiescent state in adult bone marrow (BM) (Figure S1E and S1F). Collectively, these results suggest that the eR1-CreER^T2^ mouse model is useful for HSC marking at both adult (6 weeks or older) and childhood (1-4 weeks) stages.

### eR1-CreER^T2^ system enables the best HSC enrichment

The most rigorous way to functionally define HSC is to perform transplantation assays that allow for the assessment of long-term multilineage engraftment. To further confirm HSC-specific targeting in eR1-CreER^T2^ Tg mice, long-term reconstitution experiments were performed, using the congenic Ly5.1/Ly5.2 system, where irradiated adult mice were used as recipients. Sorted BM cell suspensions were obtained from individual adult eR1-CreER^T2^ Tg; Rosa26-LSL-tdTomato mice at 48 hours after tamoxifen injection (Figure 2A). Two different sets of transplantation experiments were carried out, using either c-Kit^+^ Sca1^+^ Lin^-^ (KSL) or CD34^-^Flt3^-^KSL BM cells as donor cells. In both experiments, tdTomato^+^ and tdTomato^-^ cells were sorted from each fraction (Figure 2B).

**Figure 2.**
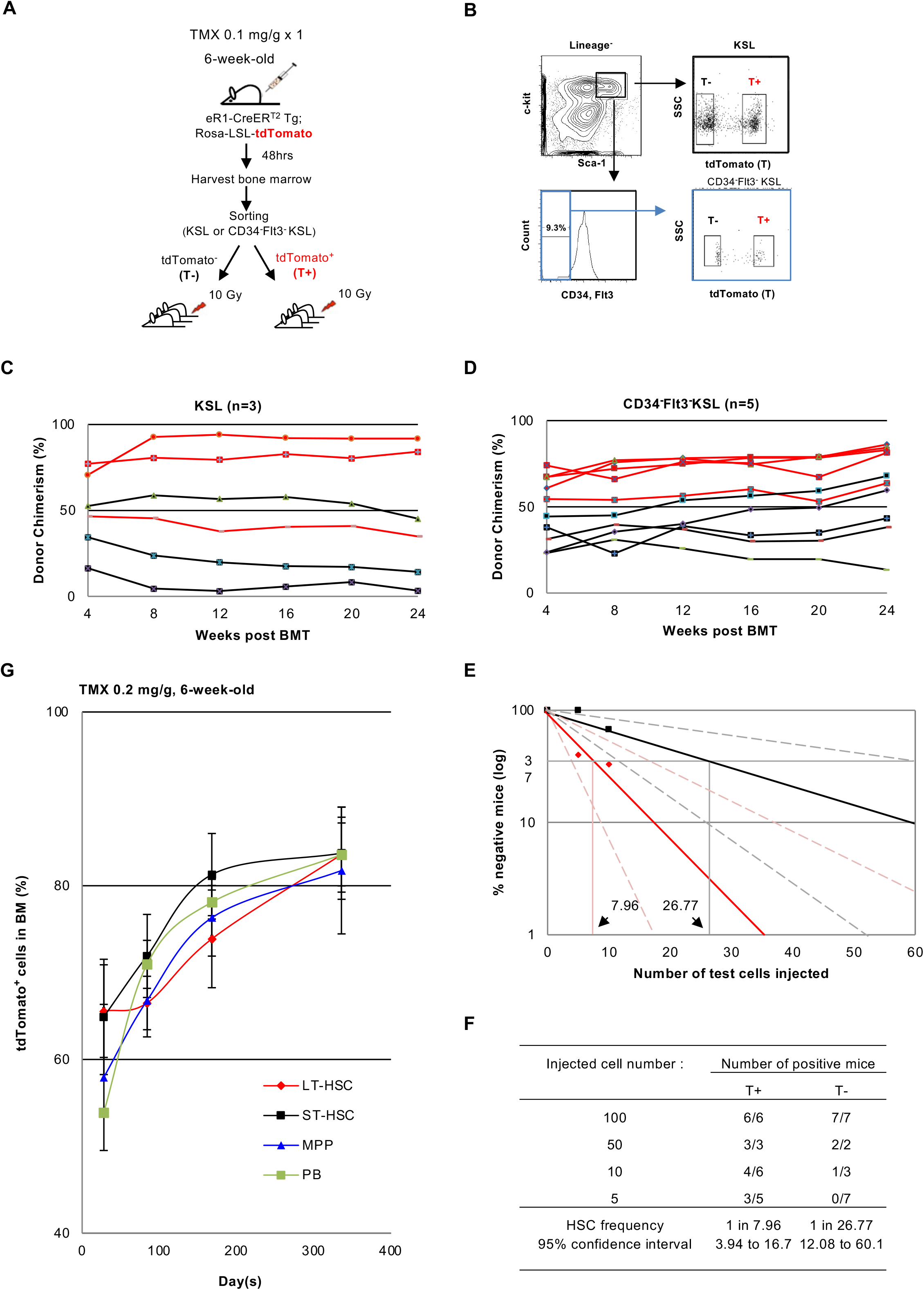
eR1-CreER^T2^ system allows for further HSC enrichment in LT-HSCs. (A) Schematic diagram depicts BMT procedure to examine eR1 activity-based enrichment of HSCs. (B) Gating strategy to isolate KSL T^+^, KSL T^-^, CD34^-^Flt3^-^KSL T^+^ and CD34^-^Flt3^-^KSL T^-^ populations. (C) & (D) Chimerism in PB of recipient mice transplanted with tdTomato^+^ (red) or tdTomato^-^ (black) cells in KSL or CD34^-^Flt3^-^KSL fractions at different time points. Each line indicates percentage of chimerism in an individual recipient mouse. (E) Limiting dilution analysis to calculate HSC frequencies in tdTomato^+^ population (red diamonds) or tdTomato^-^ population (black squares) within CD34^-^Flt3^-^KSL (LT-HSC) fraction at six months after BMT. Mice were considered negative when chimerism in PB was less than 0.5% in any examined lineage. (F) HSC frequencies were calculated using Poisson statistic with extreme limiting dilution analysis (ELDA) [30]. (G) Long-term marking in different stem cell and progenitor fractions in the bone marrow or peripheral blood from 6-week-old cohort of eR1-CreER^T2^ Tg; Rosa26-LSL-tdTomato mice. Abbreviations: T, tdTomato; PB, peripheral blood; BM, bone marrow; BMT, bone marrow transplantation; HSC, hematopoietic stem cells; LT-HSC, long term hematopoietic stem cell; ST-HSC, short-term hematopoietic stem cell.

1,000 KSL or 100 CD34^-^Flt3^-^KSL BM cells of either tdTomato^+^ or tdTomato^-^ were transplanted into irradiated recipient mice, with 2 × 10^5^ non-Tg BM cells to provide short-term radioprotection. Donor cell engraftment was examined for 6 months after transplantation. tdTomato^+^ cells from both KSL and CD34^-^Flt3^-^ KSL fractions exhibited higher short-term (1-3 months) and long-term (4-6 months) chimerism in PB respectively as compared to their tdTomato^-^ counterparts (Figure 2C and 2D).

To ascertain the HSC enrichment by eR1, limiting dilution assay was conducted. The HSC frequency of tdTomato^+^ and tdTomato^-^ populations in CD34^-^Flt3^-^KSL fraction was 1 in 7.96 and 1 in 26.77 respectively (Figure 2E and 2F). This result suggests that eR1-CreER^T2^ mouse enables the robust HSC enrichment better than the CD34^-^Flt3^-^KSL fraction which is the current standard method to sort LT-HSCs. No bias towards specific lineages in reconstitution capability was seen in eR1-active (tdTomato^+^) CD34^-^Flt3^-^KSL HSCs, whereas a moderate lymphoid-biased reconstitution was found in eR1-inactive (tdTomato^-^) CD34^-^Flt3^-^ KSL HSCs (Figure S2).

To determine the behavior of tdTomato^+^ cells in the different bone marrow stem progenitor cell fractions and peripheral blood of eR1-CreER^T2^ Tg;Rosa26-LSL-tdTomato mice, a one-year kinetic study was carried out. Similar percentages of tdTomato^+^ cells were observed in the ST-HSCs and LT-HSCs one month after tamoxifen injection (0.2 mg/g) in 6-week-old mice (Fig 2G). tdTomato^+^ cells in the ST-HSCs increased steadily at 3 months and 6 months whereas the LT-HSC tdTomato^+^ cells exhibited a slower increase than those in ST-HSCs during the same time period. However, tdTomato^+^ cells in the LT-HSC fraction increased and reached equivalent levels with ST-HSC tdTomato^+^ cells at the end of one year. These results support that eR1-mediated labeling occurs in dormant LT-HSCs, the contribution of which to native hematopoiesis is limited when young, but progressively increases with age.

### *In situ* identification of HSCs using eR1-CreER^T2^ Tg mouse

To explore the potential of eR1 system for *in situ* visualization of HSCs, bones of eR1-EGFP Tg mouse were subjected to immunohistological analysis. Half of the eR1-driven EGFP^high^ cells were c-Kit^+^ cells, and these c-Kit^+^EGFP^high^ cells, which are likely to represent HSCs, were evenly distributed along the bone of 3-week-old eR1-EGFP Tg mouse (Figure 3A arrow and 3B). In contrast, in 3-month-old mice, c-Kit^+^EGFP^high^ cells were preferentially located at the trabecular region, which provides HSCs with an environment to preserve their dormant status (Figure 3C arrow and D). This difference in the distribution of c-Kit^+^EGFP^high^ cells, consistent with the results in earlier studies, suggests that eR1-driven fluorescence signal is useful for *in situ* visualization of HSCs. A similar assay was carried out using bones from 4-month-old eR1-CreER^T2^ Tg; Rosa26-LSL-tdTomato mouse at 48 hours after tamoxifen injection. The immunoflourescent staining result showed the successful identification of c-Kit^+^tdTomato^+^ cells both in metaphysis (Figure 3E and 3G) and diaphysis (Figure 3F). In trabecular region, a c-Kit^+^tdTomato^-^ cell adjacent to a c-Kit^+^tdTomato^+^ cell was observed. Such c-Kit^+^tdTomato^-^ cell might represent a daughter cell divided from c-Kit^+^tdTomato^+^ cell by asymmetric cell division (Figure 3H). Paired c-Kit^+^tdTomato^+^ cells, which might be generated by symmetric division, were also found (Figure 3I). These results illustrate that both Tg mouse models, eR1-EGFP Tg and eR1-CreER^T2^ Tg mice, enable *in situ* imaging of HSCs.

**Figure 3.**
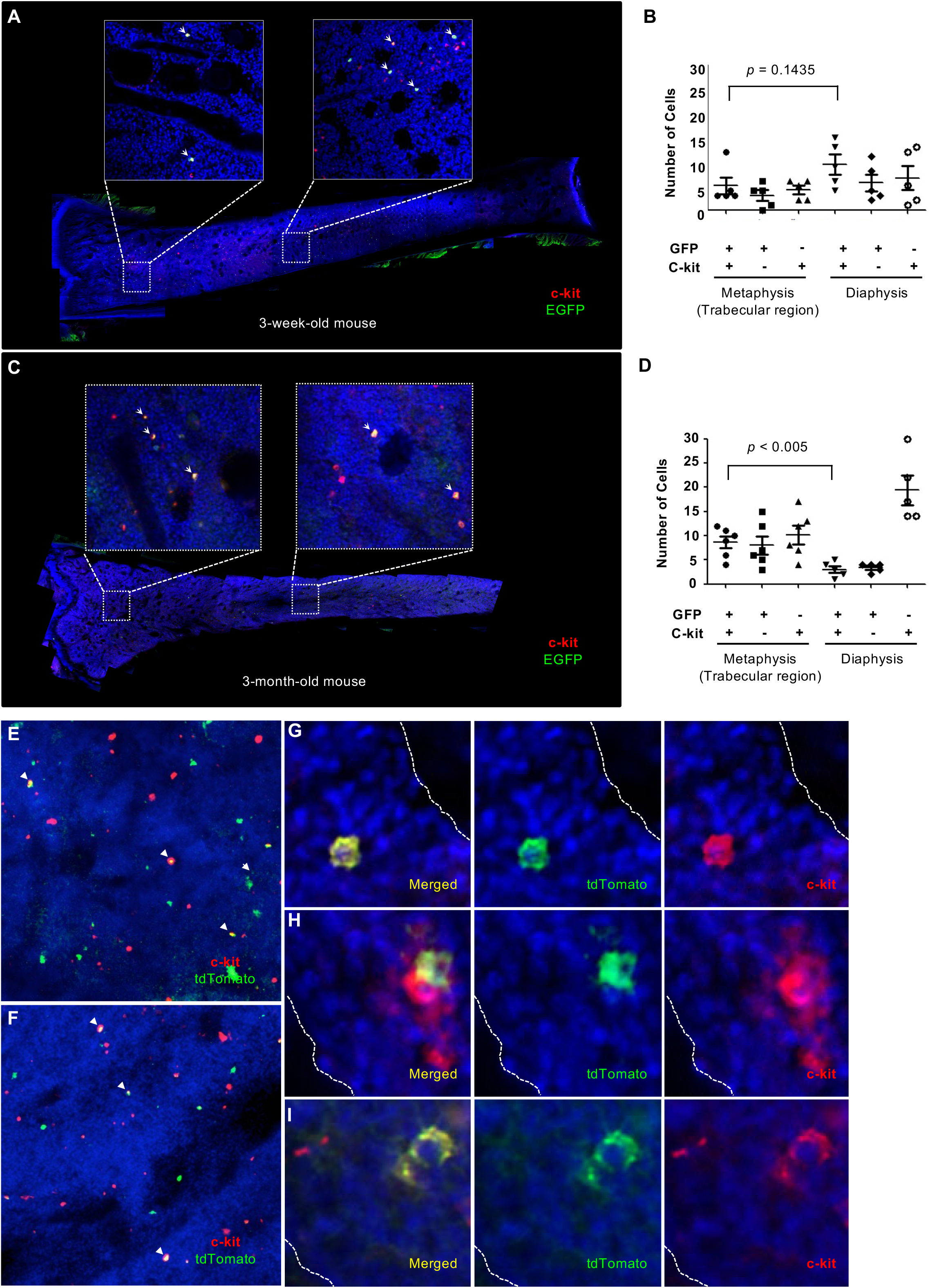
*in situ* visualization of HSCs in child and adult mouse bone marrow. (A-D) Bones from 3-week-old (A) or 3-month-old (C) eR1-EGFP Tg mice were obtained for immunostaining. Results shown in (B) and (D) were obtained from quantification of cells in (A) and (C), respectively. P-value was obtained from unpaired t test. At least five fields from each region were counted. GFP^+^c-Kit^+^ HSCs were distributed equally throughout the whole BM in 3-week-old mouse while GFP^+^ c-Kit^+^ HSCs were localized at metaphysis (trabecular region) with greater abundance compared to diaphysis in 3-month-old mouse. (E-I) Single dose 0.2 mg/g of TMX were administered into 4-month-old eR1-CreER^T2^ Tg; Rosa26-LSL-tdTomato mice. Bones were harvested 48 hours after TMX injection and subjected to immunostaining. Images were taken from (E) trabecular region and (F) diaphysis. (G), (H), and (I) are enlarged images taken from metaphysis (trabecular region). Abbreviations: TMX, tamoxifen; HSCs, hematopoietic stem cells; GFP, green fluorescence protein; EGFP, enhanced green fluorescence protein.

### Fetal HSCs are marked in eR1-CreER^T2^ Tg mouse

eR1 is shown to be active in hemogenic ECs and HPCs at embryonic stage E10.5 [2]. To investigate the eR1-driven CreER^T2^ expression pattern in newly generated Tg mouse at embryonic stage, Rosa26-LSL-tdTomato homozygous female mouse was crossed with eR1-CreER^T2^ Tg male mouse. The pregnant females were injected with 0.05 mg/g of tamoxifen on 9.5/ 10.5-and 0.05 or 0.1 mg/g 14.5-days post coitum (d.p.c.) (Figure 4A). These tamoxifen injections generated 96.7%, 87.6% and 73.2% tdTomato^+^ cells, respectively, in the PB of 3-week-old progeny mice (Figure 4B). The percentages of marked cells in PB did not increase with age of mice, unlike those observed in the adult mice. Analysis of BM cells from 6-month-old progeny mice with tamoxifen injection on 14.5-d.p.c. showed nearly 100% marking in all the hematopoietic stem/ progenitor cell (HSPC) fractions (Figure S3C).

**Figure 4.**
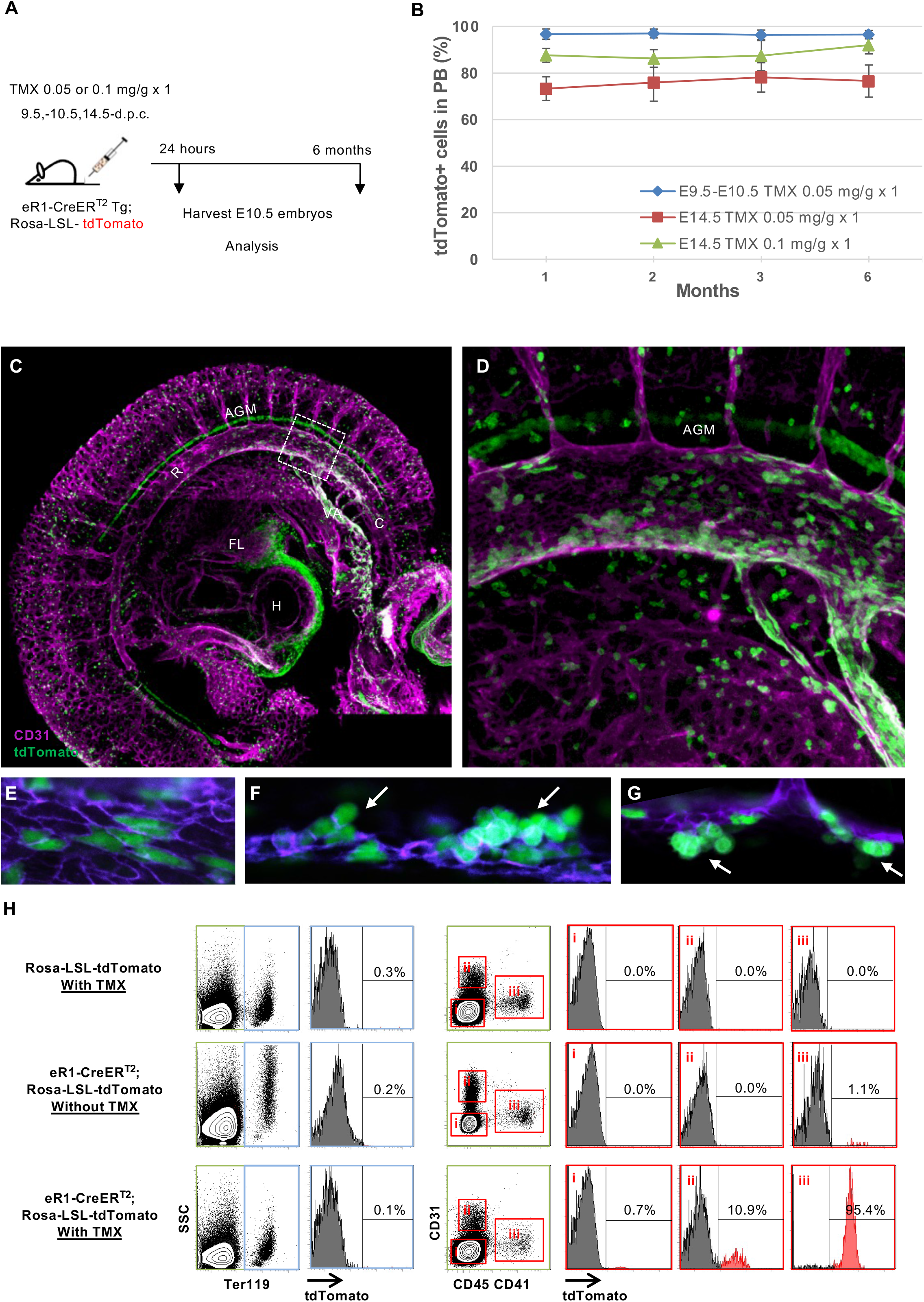
Fetal HSCs and hemogenic ECs are marked in eR1-driven CreER^T2^ Tg mice. (A) Schematic diagram demonstrates activation of eR1-driven CreER^T2^ at embryonic stage. (B) Long-term HPCs marking in PB of eR1-CreER^T2^ Tg; Rosa26-LSL-tdTomato progeny mice. The progeny mice were delivered from pregnant mothers injected with TMX on 9.5-10.5 or 14.5-d.p.c. (C) Whole-mount immunostaining of eR1-CreER^T2^ Tg; Rosa26-LSL-tdTomato E10.5 embryo with anti-CD31 (magenta) and anti-tdTomato (green) antibodies at 24 hours after TMX injection. Three-dimensional reconstruction of the eR1-CreER^T2^ Tg; Rosa26-LSL-tdTomato mouse embryo (lateral view). tdTomato signals represent CreER^T2^ activity. Within DA, the rostral (R) and caudal (C) regions are indicated. (D) Image enlarged from boxed area in (C). eR1-driven CreER^T2^ expression pattern in middle of DA within AGM region was observed. (E) Image shows ECs of DA. Some ECs were stained with tdTomato antibody. Clustered HPCs emerge from ventral (F) and dorsal (G) artery wall in AGM region. Arrows depict newly generated HPC clusters from ventral and dorsal artery wall. (H) Flow cytometry plots display the percentage of tdTomato^+^ cells in eR1-CreER^T2^ Tg; Rosa26-LSL-tdTomato E10.5 embryos. Control mice are eR1-CreER^T2^ Tg embryo (littermate of test cohort) and none TMX injected eR1-CreER^T2^ Tg; Rosa26-LSL-tdTomato embryo. Abbreviations: TMX, tamoxifen; PB, peripheral blood; UA, umbilical artery; UV, vitelline artery; AGM, aorta-gonad-mesonephros; d.p.c., days post coitum; DA, Dorsal aorta; H, heart; FL, fetal liver, HPCs, hematopoietic cells; ECs, endothelial cells; FL, fetal liver.

To examine the initially labeled cells by tamoxifen injection on 9.5-d.p.c., eR1-CreER^T2^ Tg; Rosa26-LSL-tdTomato embryos were harvested at 24 hours after tamoxifen injection. tdTomato signals were observed in the main hematopoietic sites, such as aorta-gonad-mesonephros (AGM), vitelline artery (VA), umbilical artery (UV) and fetal liver (FL), in the E10.5 embryos (Figure S3A iii). This expression pattern is largely identical to that seen in the eR1-EGFP Tg embryos (Figure S3A iv) and endogenous Runx1+ cells visualized by staining by anti-Runx1 antibody (Figure S3D). Remarkably, no tdTomato signals were observed in any eR1-CreER^T2^ Tg; Rosa26-LSL-tdTomato E10.5 embryos without tamoxifen injection (Figure S3A i and ii). Whole-mount immunostaining of the embryos further confirmed the presence of tdTomato^+^ cells in the known hematopoietic sites (Figure 4C). HPC clusters were preferentially accumulated in the middle of distal artery (DA) within the AGM region as compared to rostral and caudal regions (Figures 4C and D). HPCs at E10.5 were derived from hemogenic ECs in DA. Some endothelial cell marker CD31^+^ ECs in DA were co-stained with tdTomato antibody, suggesting that hemogenic ECs were labeled as expected (Figure 4E). tdTomato-expressing HPC clusters retained CD31 signals, indicating that the cluster forming cells were emerged from both dorsal and ventral artery walls (Figures 4F and G). The ECs of heart also displayed strong tdTomato signals. In addition, tdTomato^+^ cells were aligned between pairs of somites that most likely represent the notochord. The notochord marking was only observed in eR1-CreER^T2^ Tg mouse, but not in eR1-EGFP Tg mouse. The significance of this labeling awaits further investigation.

Labeling and gene targeting efficiency in eR1-CreER^T2^ Tg; Rosa26-LSL-tdTomato embryo was further analyzed by flow cytometry using antibodies against CD31, CD41, CD45 (pan-leukocyte marker) and Ter119 (erythrocyte marker). Consistent with the whole-mount immunostaining result, essentially all of the CD45^+^CD41^+^ cells were tdTomato^+^ cells, 11% of CD31^+^ECs exhibited tdTomato signal but Ter119^+^ erythroid cells lacked tdTomato signal (Figure 4H), suggesting that eR1 drives CreER^T2^ expression in a part of ECs, probably hemogenic ECs and HPCs, but not in primitive erythrocytes. Collectively, these results demonstrate that the eR1-CrER^T2^ Tg system is similarly useful for the labeling of hemogenic ECs and fetal HSCs.

### Induction of hematological malignancy by eR1-CreER^T2^ Tg system

It is well documented that the introduction of an oncogenic event in the stem cell fraction can efficiently induce tumor development [14-17]. To establish an inducible leukemia mouse model, eR1-CreER^T2^ Tg; Kras-LSL-G12D mice were generated. Line #3-17 of eR1-CreER^T2^ Tg mouse was utilized for this study since this Tg line demonstrated no leakage of CreER^T2^ activity in the BM of 5-month-old mouse (Figure S1D). Oncogenic Kras activation is shown to readily induce hematopoietic malignancy [14]. As expected, all eR1-CreER^T2^ Tg; Kras-LSL-G12D mice developed hematopoietic malignancies within 4 months (median survival 98 days) after a single tamoxifen injection, whereas no disease development was observed in uninjected eR1-CreER^T2^ Tg; Kras-LSL-G12D mice (Figure 5A).

**Figure 5.**
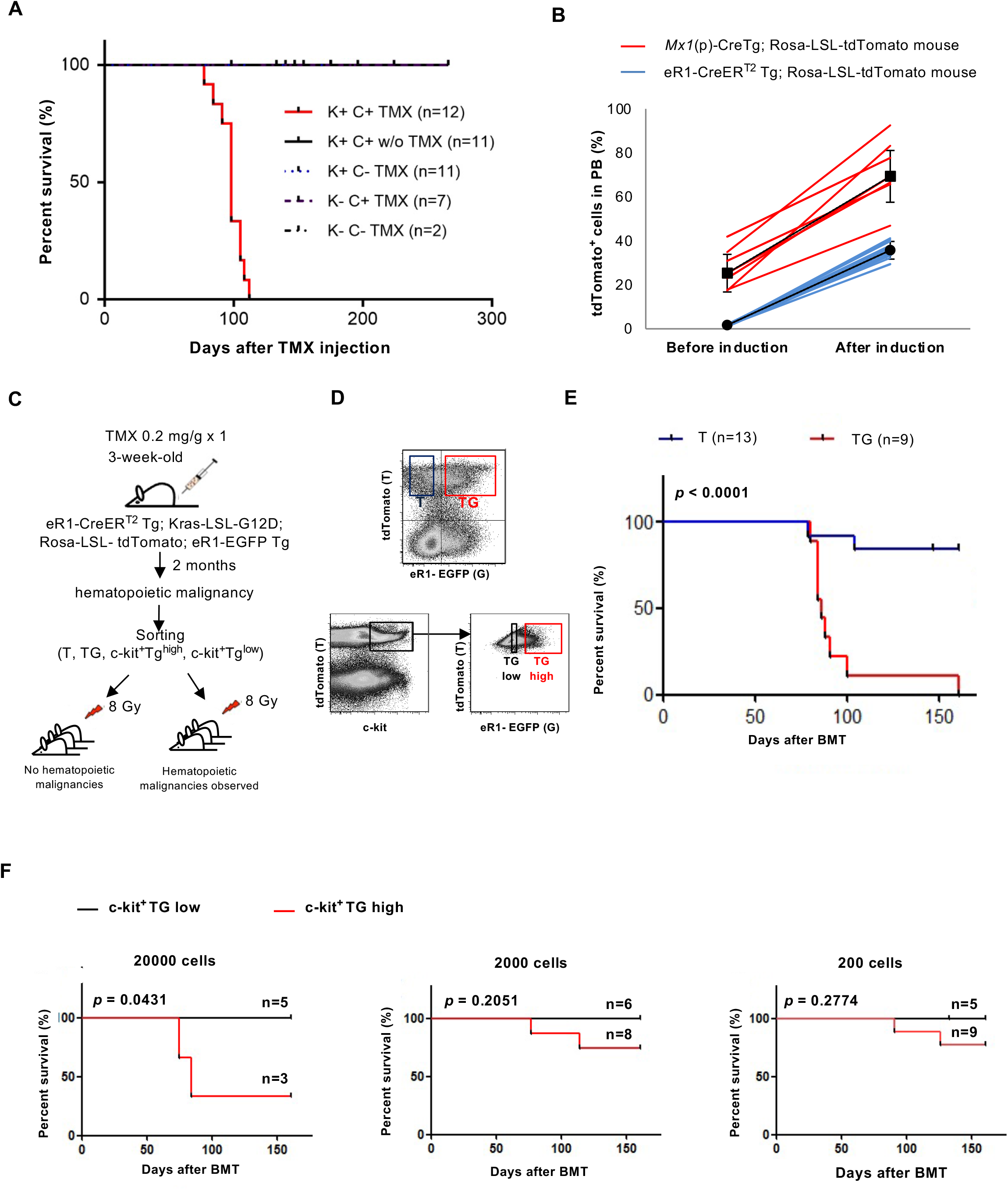
eR1 is active in leukemia stem cells. (A) Kaplan-Meier survival curve for eR1-CreER^T2^ Tg; Kras-LSL-G12D mice after TMX injection. P-value obtained from Log-rank (Mantel-Cox) Test. (B) Comparison of Cre leakage and induction efficiency between *Mx1*(p)-Cre Tg and eR1-CreER^T2^ Tg. Induction was performed in adult mice (6-week-old). For *Mx1*(p)-Cre Tg mice, 250 µg of pIpC was administered inperitoneally every alternate day for a total of 3 doses. eR1-CreER^T2^ Tg mice were injected with a single dose of TMX (0.2 mg/g) Each line represents percentage of tdTomato^+^ cells in PB of individual mouse at 1 week after induction. (C) Schematic diagram depicts BMT procedure to evaluate eR1 activity in leukemia initiating cells. BM cells were harvested from diseased mice and sorted out for BMT. (D) Gating strategy to isolate T, TG, c-Kit^+^TG^high^ and c-Kit^+^TG^low^ population. (E) Survival curve for mice transplanted with T and TG population. (F) Survival curve for mice transplanted with different amount of c-Kit^+^TG^high^ and c-Kit^+^TG^low^ population. Abbreviations: HSC, hematopoietic stem cell; TMX, tamoxifen; PB, peripheral blood; BM, bone marrow; IP, intraperitoneally; BMT, bone marrow transplantation; K, Kras-LSL-G12D; C, eR1-CreER^T2^; T, tdTomato^+^; TG, tdTomato^+^EGFP^+^; TG high, c-Kit^+^tdTomato^+^EGFP^high^; TG low, c-Kit^+^tdTomato^+^EGFP^low^.

In *Mx1* promoter (p)-Cre Tg mice, minimal Cre expression was reported to be around 5% before polyinosinic-polycytidylic acid (pIpC) injection [14]. However, this estimation was obtained using Rosa26-LSL-LacZ reporter mouse, which is far less sensitive than Rosa26-LSL-tdTomato mouse as a reporter strain. As shown in Figure 5B, *Mx1*(p)-Cre Tg; Rosa26-LSL-tdTomato mice exhibited around 20% tdTomato^+^ cells in PB of 6-week-old mice without pIpC injection and the leakage gradually increased with age (data not shown). After pIpC injection, the frequency of tdTomato^+^ cells in PB varied amongst individual mice. In contrast, eR1-CreER^T2^ Tg mice showed no leakage of Cre activity before induction and minimal variation in the percentage of tdTomato^+^ cells in PB after induction. This result reveals that the *Mx1*(p)-Cre inducible system, currently the most widely used method for the induction of leukemia, has fundamental flaws due to its high leakage of Cre activity, rendering it a faulty inducible system. Therefore, the eR1-CreER^T2^ Tg mouse is the long-awaited improved inducible mouse model for the induction of malignancy in hematopoiesis

### eR1 is active in leukemia stem cells

Cancer has been shown to harbor cells with stem cell like properties, termed cancer stem cells (CSCs) [18-20]. eR1 is anticipated to be active in CSCs in leukemia, namely leukemia stem cells (LSCs). To investigate the potential of LSC marking by eR1, tamoxifen-inducible eR1-CreER^T2^ Tg; Kras-LSL-G12D; Rosa26-LSL-tdTomato; eR1-EGFP Tg mice (henceforth referred to as 4G mice) were generated. The EGFP^+^ signal mark cells carrying stemness, while tdTomato^+^ signal serves as a surrogate marker for oncogenic Kras activation. The compound 4G mice developed hematopoietic malignancy after tamoxifen injection. BM cells were harvested from the diseased mice. EGFP^+^ and EGFP^-^ cells were separately sorted from the tdTomato^+^ population and transplanted into recipient mice (Figure 5C). Hematopoietic malignancies were observed in mice transplanted with tdTomato^+^EGFP^+^ (TG) cells, but not in those transplanted with tdTomato^+^EGFP^-^ (T) cells (Figure 5E), suggesting that eR1 is active in LSCs.

In attempt to further enrich LSCs, we employed a combination of c-Kit and EGFP intensity. A well-known HSC marker, c-Kit is often used to enrich LSCs [21]. In normal HSCs, eR1-driven EGFP^high^ fraction was found to contain greater abundance of LT-HSCs compared with eR1-driven EGFP^low^ population (Ng et al., 2010). c-Kit^+^tdTomato^+^EGFP^low^ (TG low) and c-Kit^+^tdTomato^+^EGFP^high^ (TG high) population, which was present at 1.2% in the BM of diseased 4G mouse, were isolated from diseased mice and transplanted into recipient mice (Figure 5D). In order to examine the minimal number of LSCs required to cause hematopoietic malignancies, varying number of sorted cells were used for transplantation. At all cell dosages tested, hematopoietic malignancies were observed in those transplanted with TG high population, but not in those mice transplanted with TG low population (Figure 5F). Recipient mice transplanted with as low as 200 TG high cells were capable of developing hematopoietic malignancies, suggesting that LSCs are highly enriched within c-Kit^+^tdTomato^+^EGFP^high^ population in the 4G mouse model.

### Induction of Kras^G12D^ in fetal, child and adult HSCs causes different malignancies

It has long been suspected through clinical observations that oncogenic potential and/or tumor spectrum of a certain oncogenic event is heavily dependent on the developmental stage in which it occurs. To experimentally interrogate this notion, we utilized the eR1-CreER^T2^ system to induce oncogenic Kras in HSCs at different developmental stages, such as fetus, child and adult. The results revealed that the survival curves and tumor spectrums of eR1-CreER^T2^ Tg; Kras-LSL-G12D diseased mice were distinct amongst the three cohorts (Figure 6A and 6B). In child (3-week-old) induction cohort, 67% of diseased mice were biphenotypic lymphoma carrying a mixture of myeloid and T cell features. The remaining 22% of diseased mice also exhibited myeloid features as myeloproliferative disorder (MPD) or biphenotypic leukemia. In the biphenotypic lymphomas, abnormal cells expressed myeloid markers such as Mac-1 and CD61 in thymus and spleen (Figure 6C and 6D) and immature cell marker, c-Kit (Figure 6D). In striking contrast, 100% of diseased mice developed T-cell malignancies in the adult induction cohort, including 60% lymphoma and 40% leukemia without myeloid features. Within the fetal cohort, majority of the mice developed T cell lymphoma (74%). The rest of the diseased mice (26%) displayed features of MPD and biphenotypic lymphoma with T and myeloid cell features. It was also observed that the surface marker expression of TCRαβ was higher in the T cell lymphoma of fetal cohort versus the adult cohort suggesting that the T cell lymphoma of these cohorts differ from one another (data not shown). Evidently, the induction of the same driver oncogene at distinct developmental stages resulted in remarkably different disease spectra.

**Figure 6.**
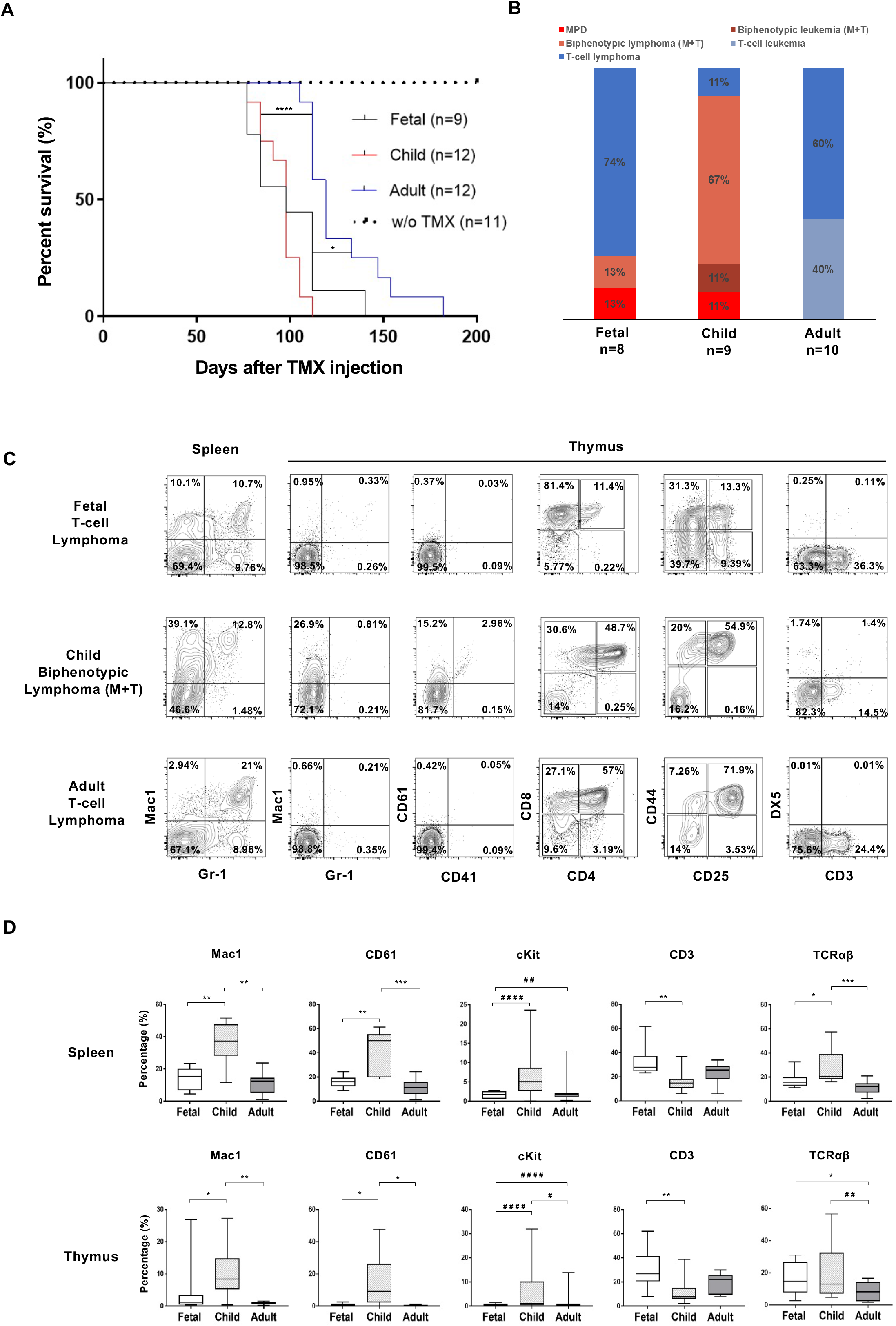
Activation of Kras^G12D^ at distinct ages demonstrates different tumor spectra. (A) Survival curve for eR1-CreER^T2^ Tg; Kras-LSL-G12D upon TMX injection at different ages. Log-rank (Mantel-Cox) test was used for calculating statistical significance. (B) Comparison of tumor spectra caused by activation of oncogenic Kras at different ages. (C) Representative flow cytometry results of diseased mice in fetal, child and adult cohorts. (D) Differences in percentage of cell surface lineage markers amongst fetal, child and adult cohorts. Asterisks or hashes represent statistical significance (*, P < 0.01; **, P < 0.001; *** P<0.0001, represents unpaired Student’s t-test; #, P<0.01; # #, P<0.001; # # #, P<0.0001, # # # #, P<0.00001, represents F test) Abbreviations: TMX, tamoxifen; w/o, without.

### A gene ablation in fetal, child and adult HSCs lead to distinct phenotypes

To demonstrate the usefulness of eR1-CreER^T2^ Tg mouse for gene targeting in HSCs, *Evi1*^fl3/fl3^ conditional knockout (cKO) mice which are known to exhibit HSC defects [22] were selected and mated to generate eR1-CreER^T2^ Tg; *Evi1*^fl3/fl3^ mice. *Evi1* is a transcription factor responsible for maintaining the integrity of HSCs. Tamoxifen was injected with a dosing regimen of 0.2 mg/g x 6 times on alternative days from 3 to 4 weeks old (childhood stage). Near complete excision was achieved in the BM at one month after tamoxifen injection (Figure 7A). Complete blood cell counts in eR1-CreER^T2^ Tg; *Evi1*^fl3/fl3^ mice exhibited thrombocytopenia and flow cytometric analysis demonstrated a decrease in LT-HSC population (Figure 7B and 7C). These phenotypes are closely resemble to those in *Evi1* cKO mice using *Mx1*(p)-Cre system, once again demonstrating that the eR1-CreER^T2^ Tg mouse is an effective new platform for common gene targeting for HSCs in hematopoiesis. Besides gene targeting in child HSCs, the *Evi1* gene was excised in fetal and adult HSCs as well. Interestingly, the phenotypes were far less pronounced. Analysis of LT-HSCs in eR1-CreER^T2^ Tg; *Evi1*^fl3/fl3^ mice among the fetal, child and adult cohorts uncovered significant differences in the LT-HSC percentages in the child cohort at the end of 1 year after tamoxifen injection, as compared to the fetal and adult cohort (Figure 7D). Concomitantly, the reduction in platelet count (thrombocytopenia) was exclusively evident in the childhood cohort in comparison to the other two cohorts (Figure 7E). This result demonstrates that the conditional deletion of *Evi1* gene at different developmental stages would give rise to divergent phenotypes.

**Figure 7.**
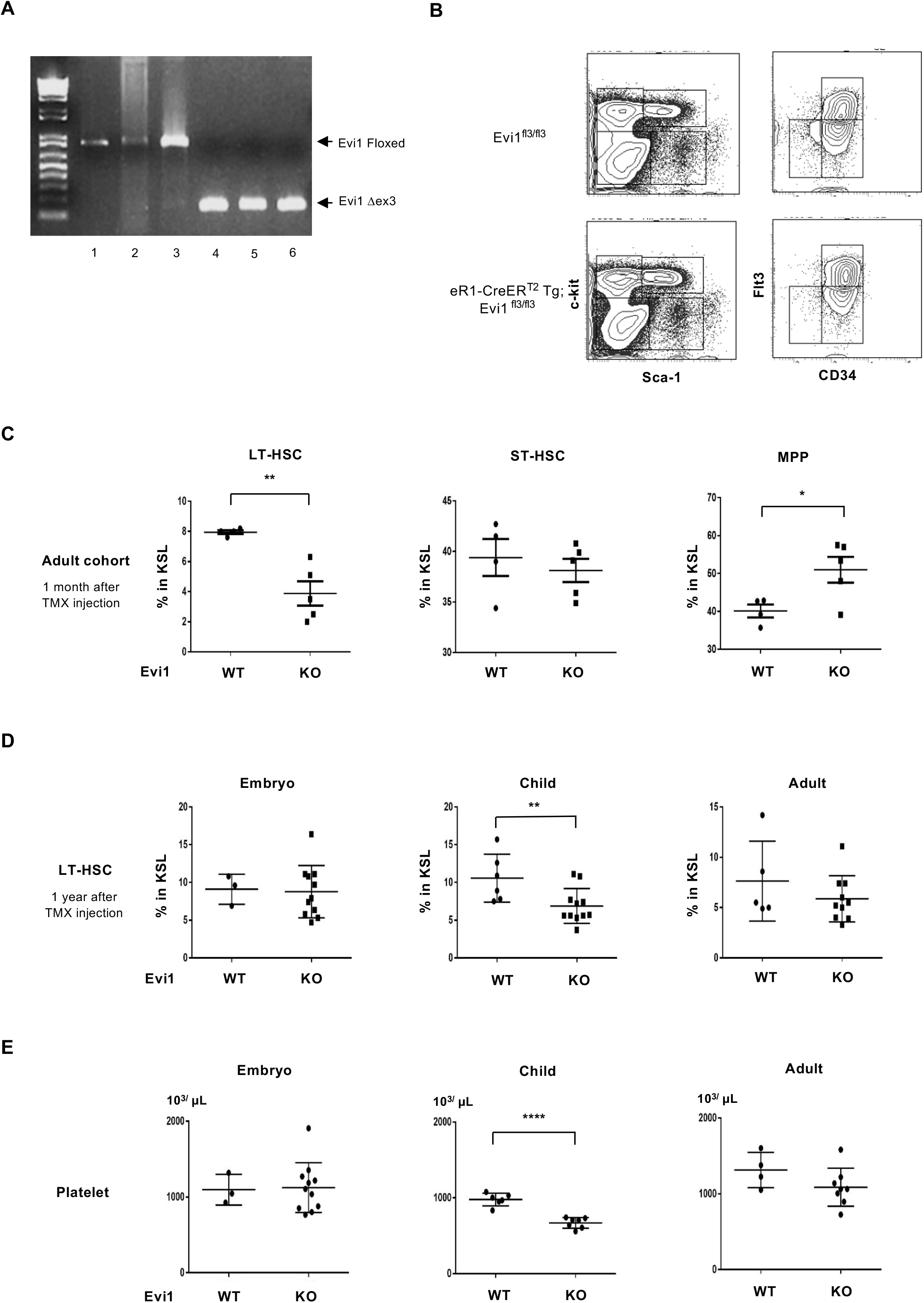
Ablation of the Evi1 gene in fetal, child and adult HSCs leads to distinct phenotypes. eR1-CreER^T2^ Tg; Evi1^fl3/fl3^mice were sacrificed one month after TMX injection. Control mice are the TMX injected littermates of eR1-CreER^T2^ Tg; Evi1^fl3/fl3^ mice without eR1-CreER^T2^ transgene. (A) Genotype PCR for Evi1 gene to detect the presence of floxed or excised exon 3. Genomic DNA from BM cells were amplified by PCR using primers (Supplemental Table 2). (B) Flow cytometric profiles of BM cells from WT and eR1-CreER^T2^ Tg; Evi1^fl3/fl3^ mice at one month after TMX injection. (C) Percentages of LT-HSC, ST-HSC and MPP in eR1-CreER^T2^ Tg; Evi1^Δ/Δ^ adult cohort mice 1 month after TMX injection. (D) Percentages of LT-HSCs in the embryo, child and adult eR1-CreER^T2^ Tg; Evi1^Δ/Δ^ mouse cohorts at 1 year after TMX injection. (E) Platelet counts in the embryo, child and adult eR1-CreER^T2^ Tg; Evi1^Δ/Δ^ mouse cohorts at 1 year after TMX injection. (*, P < 0.01; **, P < 0.001). Abbreviations: TMX, tamoxifen; BM, bone marrow; PB, peripheral blood; WT, wild-type; Evi1, Ecotropic virus integration site 1; LT-HSC, long term hematopoietic stem cell; ST-HSC, short term hematopoietic stem cell; MPP, multipotent progenitors.

## Discussion

In this report, we describe the successful generation and characterization of an inducible mouse model for highly efficient genetic engineering in HSCs, by employing the recently identified intronic enhancer for *Runx1*, eR1. The specificity and efficiency of HSC targeting achieved in this model is superior to any other mouse models currently available. A leakage of Cre activity in the absence of inducer has been a concern in the current gold standard, the *Mx1*(p)-Cre system. In clear contrast, little or no leakage occurs in our model. Besides adult HSCs, fetal and child HSCs were also readily targeted. These highly desirable attributes render the eR1-CreER^T2^ Tg mouse a powerful and compelling experimental tool for stem cell and cancer biology of the hematopoietic system.

### No leakage of Cre activity in eR1-CreER^T2^ Tg mouse model

A leakage of Cre activity in the absence of inducer has been the Achilles heel in the current benchmark methodology, *Mx1*(p)-Cre system. Crucially, the unpredictable amount of leakage is likely to compromise experimental design and interpretation of mouse phenotypes examined. The spontaneous activation of oncogenic Kras in uninduced *Mx1*(p)-Cre Tg mouse leading to the development of hematological malignancies clearly highlights this shortcoming. Whereas *Mx1*(p)-Cre Tg; Kras-LSL-G12D mice predominantly develop MPD after pIpC injection [23, 24], these mice, surprisingly, also developed MPD without pIpC injection, with 100% penetrance, with disease latency similar to that of pIpC-injected mice [24]. Therefore, the *Mx1*(p)-Cre Tg mice is not a suitable inducible system as a leukemia model. In clear contrast, the eR1-CreER^T2^ Tg mouse serves as a long-awaited leakage-free inducible system for HSC-specific genetic engineering in mice.

One caveat, however, is that the CreER^T2^ is an extremely ligand-sensitive system, where a very small amount of tamoxifen is sufficient to trigger CreER^T2^ activity. This sensitivity is particularly critical when the CreER^T2^ system is used for establishing a leukemia mouse model. We observed disease manifestation in uninjected eR1-CreER^T2^ Tg; Kras-LSL-G12D mice when housed with tamoxifen injected mice in the same cage, indicating cross-contamination via excretions of injected mice. No such diseased mice were observed when tamoxifen injected and non-injected mice were separately housed. Therefore, in order to avoid inadvertent activation of CreER^T2^ in eR1-CreER^T2^ Tg mouse, tamoxifen injected and non-injected mice should be housed in separate cages.

### Age-dependent differences in HSC property

*Vav*(p)-Cre Tg mouse and *Mx1*(p)-Cre Tg mouse are two commonly used mouse models for HSC-specific gene targeting. As *Vav*(p)*-* and *Mx1*(p)-Cre activities are both prominent in hematopoietic tissues, the major phenotypes observed, including those in the HSC populations, are largely similar when utilizing either genetic element for targeting a specific gene. However, distinct phenotypes have also been observed when the same gene was targeted by these two systems [25]. These differences could be explained by distinct requirement of a gene in developmental versus adult hematopoiesis. *Vav*(p) is activated at E10.5 onwards during the critical period of rapid HSC expansion, whereas *Mx1*(p) is induced at the adult stage when HSCs are largely in quiescence. It is therefore plausible that phenotypic descriptions about a plethora of gene targeting mice using *Mx1*(p)-Cre Tg system are a result of a mixture of developmental and adult stage dependent phenotypes. In light of this, all previously reported mouse phenotypes should be revisited. For this purpose, the eR1-CreER^T2^ Tg mouse appears to be an invaluable tool as it enables a leakage-free Cre induction at all developmental stages.

It is well documented that many types of cancer show age-dependent prevalence. For instance, trisomy 21, the most common congenital chromosomal abnormality, is at high leukemia risk in fetal and childhood hematopoiesis, but not in adult bone marrow [26]. This difference is attributed to the inherent property of background cells, particularly of the HSCs. Here, we observed that the induction of oncogenic Kras in HSCs of different developmental stages led to distinct spectra of malignancies. Interestingly, the induction at child HSCs caused leukemia/lymphomas of myeloid features, whereas the induction in fetal and adult HSCs largely induced T-cell malignancies. eR1-CreER^T2^ mouse appears to serve as a promising experimental platform to recapitulate clinical observation, namely age-dependent tumorigenesis, thereby allowing for elucidation of its underlying molecular mechanism.

As exemplified above, many scientifically important questions can be addressed using the eR1 enhancer system. Besides being a powerful tool in the HSC biology, we recently found that eR1 is also active in stem cells in stomach [27] and seemingly in multiple other organs. Extensive studies for individual tissue stem cells are currently underway. The common stemness machinery governing multiple tissue stem cells may be unveiled via the eR1 enhancer-related investigation and experimental tools, and would contribute to tissue engineering and regenerative medicine.

### Experimental Procedures

#### Mice

eR1-CreER^T2^ transgenic mice (Tg) were generated as described (supplemental information) and maintained in C57BL/6 background. Rosa26-LSL-tdTomato mice (B6;129S6-Gt (Rosa)2Sor^tm9(CAG-tdTomato)Hze^/J) were purchased from Jackson laboratory. To analyze CreER^T2^ recombinase activity, eR1-CreER^T2^ Tg mice were crossed with the reporter strain, Rosa26-LSL-tdTomato mouse. eR1-CreER^T2^ Tg; Rosa26-LSL-tdTomato mice were heterozygous at both eR1-CreER^T2^ Tg and Rosa26-LSL-tdTomato loci. eR1-CreER^T2^ Tg and Rosa26-LSL-tdTomato mice were used as controls. For the activation of CreER^T2^ recombinase, mice received intraperitoneal (IP) tamoxifen (Sigma T5648) injection at different doses for different ages as stated. Tamoxifen was dissolved in sunflower seed oil (Sigma S5007)/ethanol (10:1) mixture at 10 mg/ml. eR1-EGFP mice [2], Evi1 exon 3 floxed mice [22], Mx1-Cre [28], Kras-LSL^G12D^ [29] and CD45.1 (Ly5.1) mice are described elsewhere and all maintained in C57BL/6 background in the mouse facilities, National University of Singapore. All animal experiments followed strict guidelines set by National Advisory Committee for Laboratory Animal Research and approved by the Institutional Animal Care and Use Committee (IACUC) of National University of Singapore (NUS).

### Statistical analysis

Student’s two-tailed, unpaired t-test and F test were used to determine statistical significance of two sets of samples. Log-rank (Mantel-Cox) test was used to determine statistical significant of two survival curves. Kruskal Wallis and Mann Whitney t test was performed for comparing two non-parametric data sets. P-values less than 0.05 were considered significant. The statistical tests were performed using the Graph Pad Prism 7 (GraphPad Software Inc., La Jolla, CA)

## Supporting information

Koh et al Suppl Text Table Fig

## Acknowledgements

We thank members of MD2 Vivarium, NUS, for mouse husbandry. This work was supported by National Medical Research Council, Biomedical Research Council, A*STAR (Agency of Science, Technology and Research), the National Research Foundation Singapore, the Singapore Ministry of Education under its Research Centres of Excellence initiative, Joint NCIS and N2CR Seed Funding, Japanese Society of Hematology, Leukemia Research Foundation, and JSPS Kakenhi Grant Numbers JP18KT0026 and JP119K07668, Japan.

## Notes

### Competing Interest Statement

The authors have declared no competing interest.

