## Supplementary material for "Highly efficient *Runx1* enhancer eR1-mediated genetic engineering for fetal, child and adult hematopoietic stem cells": Koh et al Suppl Text Table Fig

**Supplemental Information**

**Figure S1. Generation of eR1-CreER$^{T2}$ Tg lines with efficient induction of Cre recombinase activity. Related to Figure 1.**

(A) Percentage of tdTomato$^+$ cells in the peripheral blood (PB) of five eR1-CreER$^{T2}$ Tg lines one week after TMX injection.

(B) Percentage of tdTomato$^+$ cells in the bone marrow (BM) of 6-week-old cohort of eR1-CreER$^{T2}$ Tg; Rosa26-LSL-tdTomato mouse, at 3 months after a single 0.2 mg/g TMX injection.

(C) Percentage of tdTomato$^+$ cells in BM of 5 months old eR1-CreER$^{T2}$ Tg line #5-2; Rosa26-LSL-tdTomato control mouse without TMX injection.

(D) Percentage of tdTomato$^+$ cells in BM of 5 months old eR1-CreER$^{T2}$ Tg line #3-17; Rosa26-LSL-tdTomato control mouse without TMX injection.

(E & F) Short-term (30 days) kinetics of tdTomato$^+$ cells in PB of eR1-CreER$^{T2}$ Tg; Rosa26-LSL-tdTomato mice with single 0.2 mg/g TMX injection into (E) 3-week-old and (F) 6-week-old mice. Mice were sacrificed at indicated time points and BM cells were harvested for flow cytometric analysis. At least three mice were analyzed in each time point for individual cohort. Mean ± S.D. of tdTomato$^+$ BM cells at each time point is shown.

Abbreviations: TMX, tamoxifen; LT-HSC, long-term HSC; ST-HSC, short-term HSC; MPP, multipotent progenitors; KSL, c-Kit+Sca-1+Lin-; KL, c-Kit+Lin-; Lin +ve, lineage markers positive.

**Figure S2. Non-biased multilineage long-term reconstitution by eR1-active (tdTomato$^+$) CD34$^-$Flt3$^-$KSL HSCs in limiting dilution transplantation assay. Related to Figure 2.**

Red lines represent chimerisms in recipient mice transplanted with eR1-active (tdTomato$^+$) CD34$^-$Flt3$^-$KSL HSCs, while black lines represent those in recipient mice transplanted with eR1-inactive (tdTomato$^-$) CD34$^-$Flt3$^-$KSL HSCs. Mean $\pm$ S.D. chimerism in the peripheral blood (PB) of recipient mice at each time point is shown in log scale.

**Figure S3. Near complete labeling of fetal HSCs in eR1-CreER$^{T2}$; Rosa26-LSL-tdTomato mice. Related to Figure 4.**

(A) E10.5 embryo images. Bright field (i) and fluorescent (ii) images of eR1-CreER$^{T2}$ Tg; Rosa26-LSL-tdTomato E10.5 embryos without tamoxifen (TMX) induction. (iii) Fluorescent image of eR1-CreER$^{T2}$ Tg; Rosa26-LSL-tdTomato E10.5 embryo with 0.05 mg/g TMX injection on day E9.5 post coitum. (iv) Fluorescent image of eR1-EGFP Tg E10.5 embryo.

(B) Percentages of tdTomato$^{+}$ cells in BM hematopoietic stem progenitor cell (HSPC) fractions of 6-month-old eR1-CreER$^{T2}$ Tg; Rosa26-LSL-tdTomato progeny mice with 0.05 mg/g TMX injection on day E9.5 post coitum.

(C) Percentages of tdTomato$^{+}$ cells in BM HSPC fractions of 6-month-old eR1-CreER$^{T2}$ Tg; Rosa26-LSL-tdTomato progeny mice with 0.1 mg/g TMX injection on day E14.5 post coitum.

(D) (i) Whole mount immunostaining of WT E10.5 embryo with anti-CD31 (magenta) and Runx1 (green) antibodies. (ii) Arrowhead showing clustered HPCs emerge from the ventral and dorsal artery wall in AGM region (image enlarged from the box area). (iii) Image shows ECs of DA. Some ECs were stained with Runx1 antibody.

Abbreviations: DA, dorsal aorta; AGM, aorta-gonad-mesonephros; HPCs, hematopoietic cells; EC, endothelial cell.

**Figure S4. Distinct tumor spectrums in fetal, child and adult cohorts in eR1-CreERT2 Tg; Kras-LSL-G12D mice. Related to Figure 6.**

(A & B) Different cell surface marker profiles analyzed in the spleen and thymus of the diseased mice from the fetal, child and adult cohort.

(C) Organ weights from the spleen, thymus and liver of the diseased mice from the fetal, child and adult cohorts.

(D) WBC, hemoglobin and platelet counts recorded from PB of the diseased mice from the three different mice age cohorts.

Abbreviations: WBC, white blood cell; PB, peripheral blood.

For T test: *, P < 0.01; **, P < 0.001; *** P<0.0001). For F test: #, P<0.01; # #, P<0.001; # # #, P<0.0001,

**Table S1: Classification of tumors in eR1-CreER$^{T2}$ Tg; Kras-LSL-G12D diseased mice**

| Group | Subtype | WBC count | Immunophenotype | Spleen weight ** | Thymus weight** |
|---|---|---|---|---|---|
| 1 | MPD | Normal/high* | Myeloid | ++ ~ +++ | - ~ + |
| 2 | Biphenotypic leukemia (M+T) | High* | Myeloid and Lymphoid (T) | ++ ~ +++ | - ~ +++ |
| 3 | Biphenotypic lymphoma (M+T) | Normal | Myeloid and Lymphoid (T) | ++ ~ +++ | - ~ +++ |
| 4 | T cell leukemia | High* | Lymphoid (T) | +++ | +++ |
| 5 | T cell lymphoma | Normal | Lymphoid (T) | ++ ~ +++ | ++ ~ +++ |

* Higher than 20 x $10^3$/μl
** + represents 150-300 mg; ++, 301-600 mg; +++, 601 mg or above.
Abbreviations: WBC, white blood cell; MPD, myeloproliferative disorder; M, myeloid; T, T-cell.

**Table S2: Mouse genotyping primer sequences**

| Gene | Forward Primer Sequence | Reverse Primer Sequence | Product size (bp) |
|---|---|---|---|
| **eR1** | CACTGATAACGTGGGCAGCTT | GTGTCCGGTGACGTGATCCTC | 400 |
| **Rosa26** | AAAGTCGCTCTGAGTTGTTAT | GGAGCGGGAGAAATGGATATG | 600 |
| **tdTomato** | GGCATTAAAGCAGCGTATCC | CTGTTCCTGTACGGCATGG | 196 |
| **Mx1-Cre** | GCGGACGGAGCACTATTTA | CCGGCATCAACGTTTTCTTT | 450 |
| **EGFP** | TGAACCGCATCGAGCTGAAGGG | TCCAGCAGGACCATGTGATCGC | 350 |
| **Kras** | CTAGCCACCATGGCTTGAGT | TCCGAATTCAGTGACTACAGATG | 400 |
| **Evi1** | CAGCTTAGACCTCAGCTAAC | GAAGAGCTCTTGCTGTTCATG | WT- 269<br>Floxed- 375 |

#### Supplemental Experimental Procedures

##### Generation of eR1-CreER$^{T2}$ transgenic mice.

eR1-CreER$^{T2}$ construct was linearized from the vector by restriction enzyme digestion, gel purified (QIAGEN, Germany), resuspended in plasmid injection buffer (10 mM Tris-HCl, 0.1 mM EDTA, pH7.4), and then sent to the Transgenic and Gene Targeting Facility at Cancer Science Institute of Singapore for the generation of transgenic (Tg) mice.

##### Analysis of eR1-CreR$^{T2}$ Tg; Kras-LSL-G12D disease mice

The spleen and thymus of eR1-CreR$^{T2}$ Tg; Kras-LSL-G12D mice from the different cohorts that developed hematological malignancies were analyzed by flow cytometry. The cells were harvested as previously described [1]. The following antibodies were used for analysis of cell surface markers: FITC conjugated antibodies against Gr1 (RB6-8C5), CD34 (RAM34), CD41 (MWReg30), CD71 (C2), CD4 (RM 4-5), TCRαβ (H57.597), CD25 (eBioCD4), B220 (RA3-6B2), CD3 (145-2C11); PE conjugated antibodies against Mac1 (M1/70), c-Kit (2B8), Fas (Fas), CD61 (CD61), Ter119 (Ter119), TCRγδ (eBioGL3), CD44 (IM7), CD19 (1D3), Dx5 (Dx5), CD8 (53-6.7). All antibodies were purchased from BD Pharmingen (San Jose, California) or eBioscience. Data were analyzed suing FowJo (v10.0) and FACS DIVA (v8.0) softwares.

##### Flow cytometric analyses and cell sorting

Peripheral blood (PB), spleen, thymus or bone marrow (BM) cells were harvested from mice and stained with respective antibodies as previously described. The following antibodies were used for identification of cell surface markers: FITC-conjugated antibodies against Gr1 (RB6-8C5), Mac-1 (M1/70), Ter119 (Ter-119), CD3 (145-2C11), CD4 (RM4-5), CD8 (53-6.7), B220 (RA3-6B2), and IL7Rα (A7R34) were used to exclude lineage positive BM cells; PE-Cy7-conjugated-anti-c-Kkit (2B8) and PerCP-Cy5.5-conjugated-anti-Sca-1 (Clone D7) antibodies were utilized for positive selection of HSPCs. tdTomato fluorescent protein was detected in PE channel. In addition, tdTomato expression is investigated in sub-fractions within the c-Kit+Sca1+Lin- (KSL) compartment, using a combination of APC conjugated CD34 (RAM34) and biotin Flt3 (A2F10) and streptavidin APC-Cy7 antibodies. All antibodies were purchased from BD Pharmingen (San Jose, California) or eBioscience. Hoechst 33258 was used for dead cell discrimination. Flow cytometric analyses and cell sorting were performed using LSR II or FACS Aria (BD Biosciences). Data were analyzed using FACS DIVA (v8.0) and FlowJo softwares (v10).

**Bone marrow transplantation**

BM cells were harvested at 48 hours after tamoxifen injection (0.1 mg/g) from eR1-CreER$^{T2}$ Tg; Rosa26-LSL-tdTomato mice (test cells, CD45.2/CD45.2) and transplanted as previously described [1]. For the mouse leukemia model, recipient mice were irradiated with a dose of 8 Gy whereas for long term-reconstitution or limiting dilution assays, recipient mice were irradiated with a dose of 10 Gy.

PB was drawn monthly from the retro-orbital plexus and subjected to flow cytometric analysis after BMT. Multilineage contribution of donor cells to hematopoiesis was assessed by analyzing the percentage of Ly5.2$^{+}$/Ly5.1$^{-}$ cells in myeloid cells, T- and B-cells respectively. The mixture of APC-conjugated-anti-Gr1 (RB6-8C5), APC-conjugated-anti-Mac-1 (M1/70), APC-conjugated-anti-B220 (RA3-6B2), PE-Cy7-conjugated-anti-B220 (RA3-6B2), PE-Cy7-conjugated-anti-CD3 (145-2C11), FITC-conjugated-anti-CD45.1 (A20) and APC-Cy7-conjugated-anti-CD45.2 (104) antibodies were utilized in the multilineage chimerism analysis. Antibodies were purchased from BD Pharmingen (San Jose, California) or eBioscience.

For limiting dilution assay, recipient mice with donor contribution lower than 0.5% in either lineage were considered as negative mice. The number of hematopoietic stem cells (HSCs) was calculated using extreme limiting dilution analysis (ELDA) [2]. The end point of the BMT experiment was six months post-BMT and multilineage chimerism analyses were performed using PB, BM, spleen and thymus cells.

**Single cell dissociation of E10.5 mouse embryo**

E10.5 embryo was suspended in 1 mL of 2 mg/mL dispase (Gibco BRL, Grand Island, NY) and incubated at 37$^{0}$C water bath for 30 minutes, with vigorous shaking occasionally during this incubation period. 1 mL of washing buffer [20% FBS; 5 mM $CaCl_2$ & Dnase 1 (50 ug/ml) in PBS] was added and the sample was centrifuged at 1,500 r.p.m. for 5 minutes at 4$^{0}$C. The supernatant was aspirated and the cell pellet was washed with 1 mL of PBS followed by another round of incubation of the cell suspension with 1 mL of Cell Dissociation Buffer (Gibco BRL) at 37$^{0}$C for 30 minutes. The cell pellet was washed and passed through 40 μm strainer to remove undissociated tissues, blocked with mouse serum and stained with respective antibodies for flow cytometric analysis.

**Whole-mount immunostaining of E10.5 embryo**

Mouse embryos from timed mating were dissected at E10.5, wherein the day of detection of the vaginal plug was designated as E0.5. Whole-mount embryo immunostaining were performed as described previously [3]. In brief, E10.5 embryo were fixed for 20-30 minutes in 2% paraformaldehyde/phosphate buffered saline (PBS) (6 ml) on ice followed by dehydrated in graded concentrations of methanol/PBS (50%, 100%). The yolk sac and rostral half of the body, from which limb buds and lateral body wall were removed, were rehydrated through 50% methanol/PBS and several ice-cold PBS washes. Embryos were blocked with PBS-MT (PBS containing 1% skim milk and 0.4% Triton-X100) followed by overnight primary and secondary antibodies incubation. Primary antibodies were biotinylated rat anti-mouse CD31 (MEC13.3, BD Pharmingen, 1/500 dilution), rabbit anti-EGFP (MBL 598, 1/2000 dilution) and rabbit anti-RFP (MBL PM005, 1/1000 dilution). Secondary antibodies were goat anti-rat IgG-Alexa 555 (Invitrogen) and goat anti-rabbit IgG conjugated to Alexa 647 (Invitrogen). Immuno-stained embryos were dehydrated and mounted in a 1:2 mix of benzyl alcohol and benzyl benzoate (BABB) to increase the transparency of tissues, followed by imaging with a confocal microscope (Zeiss LSM510). Three-dimensional reconstructions were generated from z-stacks with LSM Image Brows (Zeiss).

**Analyses of peripheral blood (PB) cells**. PB was collected by retro-orbital bleeding and complete blood cell count (CBC) was performed using automated hematology analyzer (Hemavet 950 FS, Drew Scientific or Celltac alpha MEK-6358, Nihon Kohden Corp., Tokyo, Japan).

**Genotyping.** Genomic DNA was extracted from mouse tails using Direct PCR Lysis Reagents (Viagen Biotech, Los Angeles, CA). Genomic DNA samples were then used to genotype for the respective genes by PCR. The gene specific PCR primers are listed in Table S2.

### Figure S1

A

| Line # | tdTomato⁺ cells | |
|---|---|---|
| | Before TMX injection | 1 week after TMX injection |
| 3-13 | 0.25% | 5.45% |
| 3-17 | 0.01% | 1.47% |
| 4-26 | 0.11% | 1.98% |
| 5-2 | 0.82% | 35.83% |
| 5-5 | 0.03% | 2.09% |

B

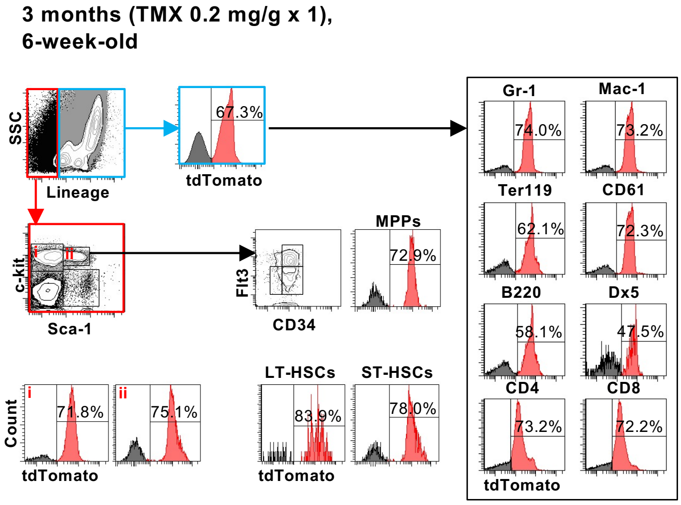


C

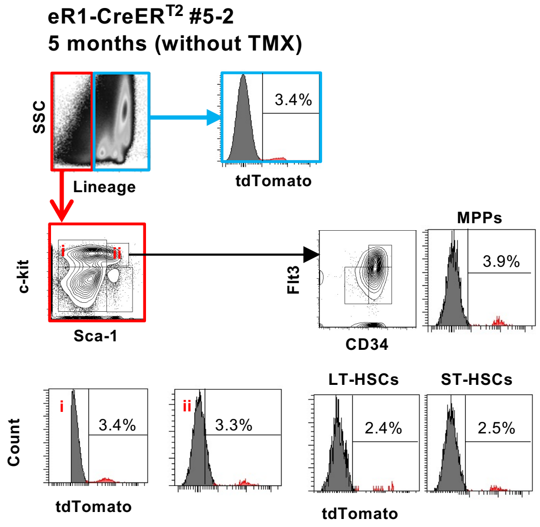


D

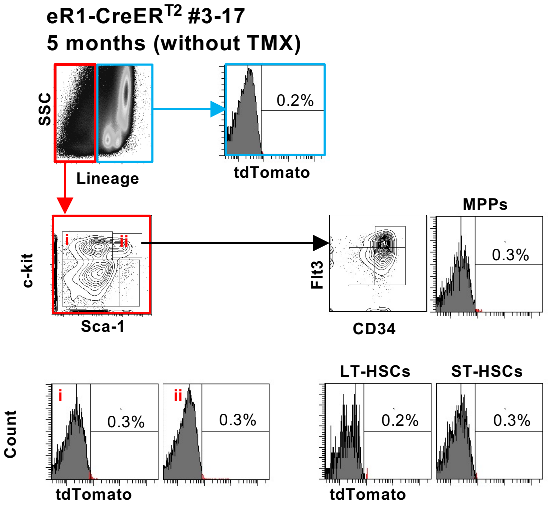


E

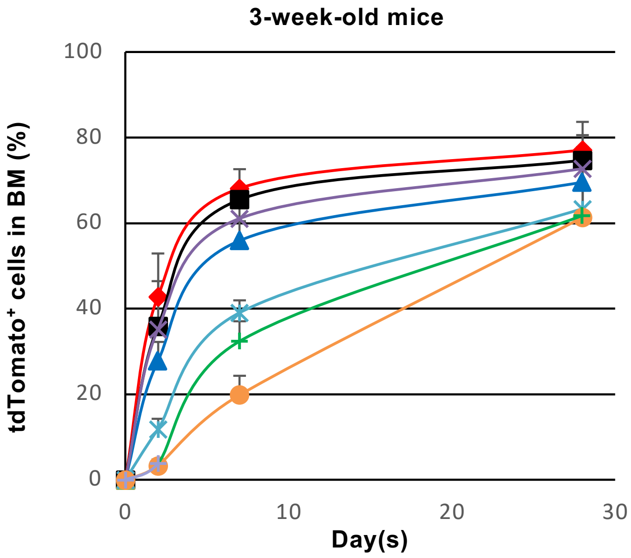


F

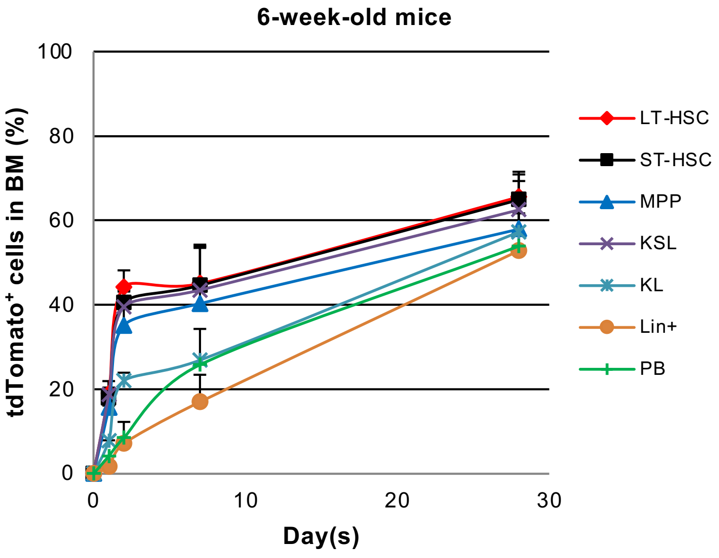

### Figure S2

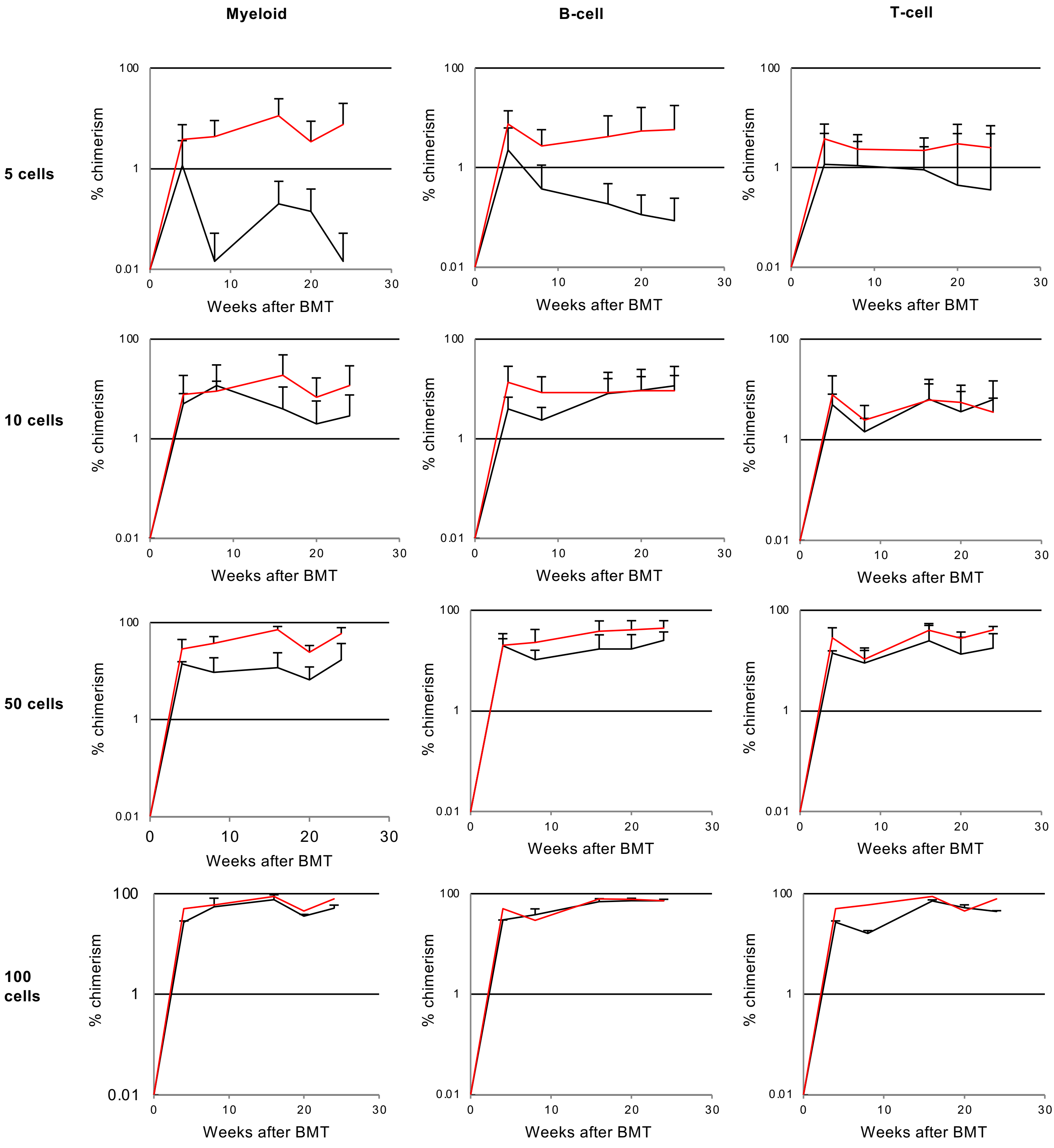
Myeloid
B-cell
T-cell
5 cells
10 cells
50 cells
100 cells
% chimerism
100
1
0.01
0
10
20
30
Weeks after BMT

### Figure S3

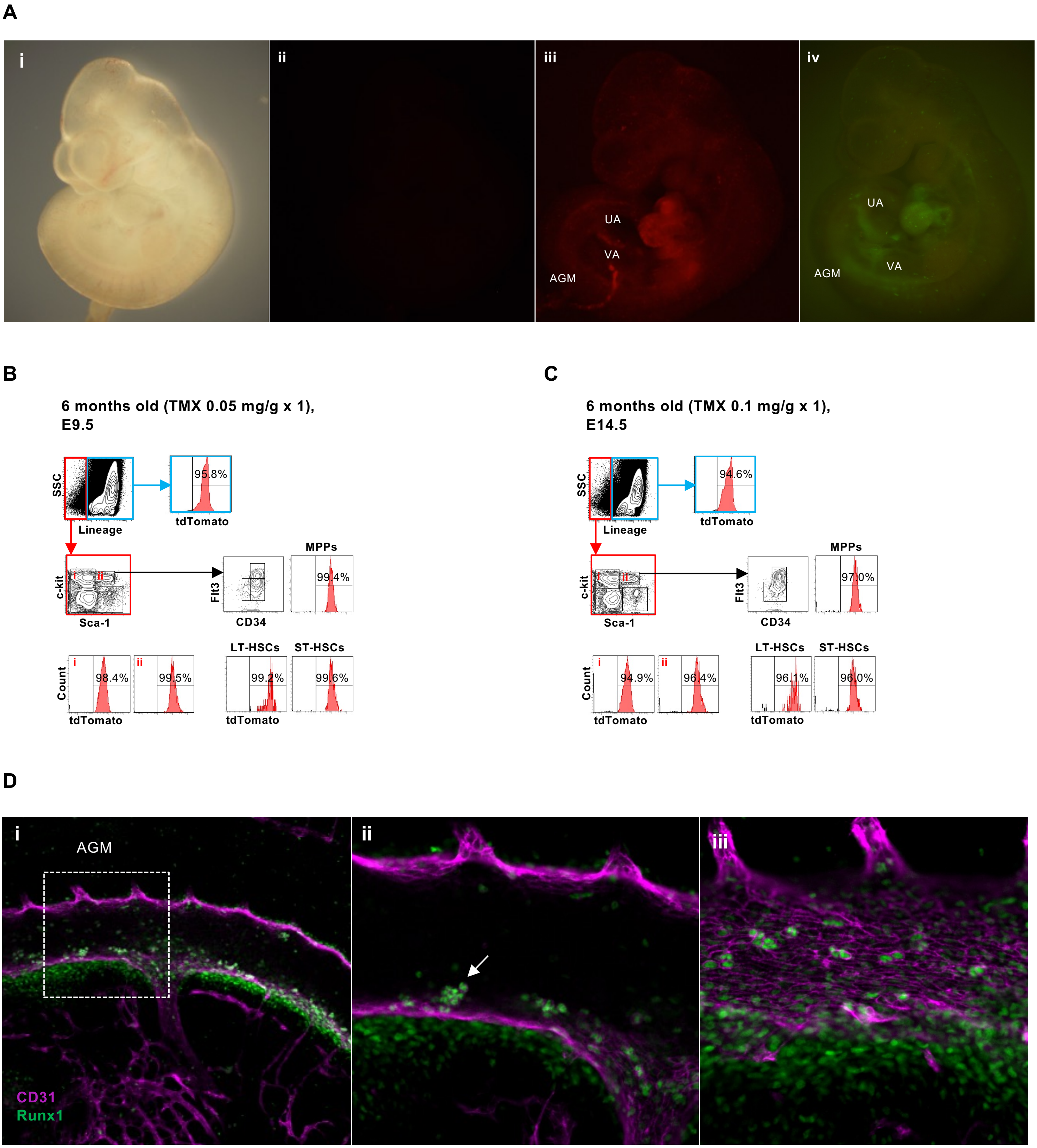

A
i
ii
iii
iv
UA
VA
AGM
UA
AGM
VA
B
6 months old (TMX 0.05 mg/g x 1),
E9.5
SSC
Lineage
95.8%
tdTomato
c-kit
Sca-1
Flt3
CD34
MPPs
99.4%
Count
98.4%
99.5%
tdTomato
LT-HSCs
ST-HSCs
99.2%
99.6%
tdTomato
C
6 months old (TMX 0.1 mg/g x 1),
E14.5
SSC
Lineage
94.6%
tdTomato
c-kit
Sca-1
Flt3
CD34
MPPs
97.0%
Count
94.9%
96.4%
tdTomato
LT-HSCs
ST-HSCs
96.1%
96.0%
tdTomato
D
i
AGM
CD31
Runx1
ii
iii

#### Figure S4

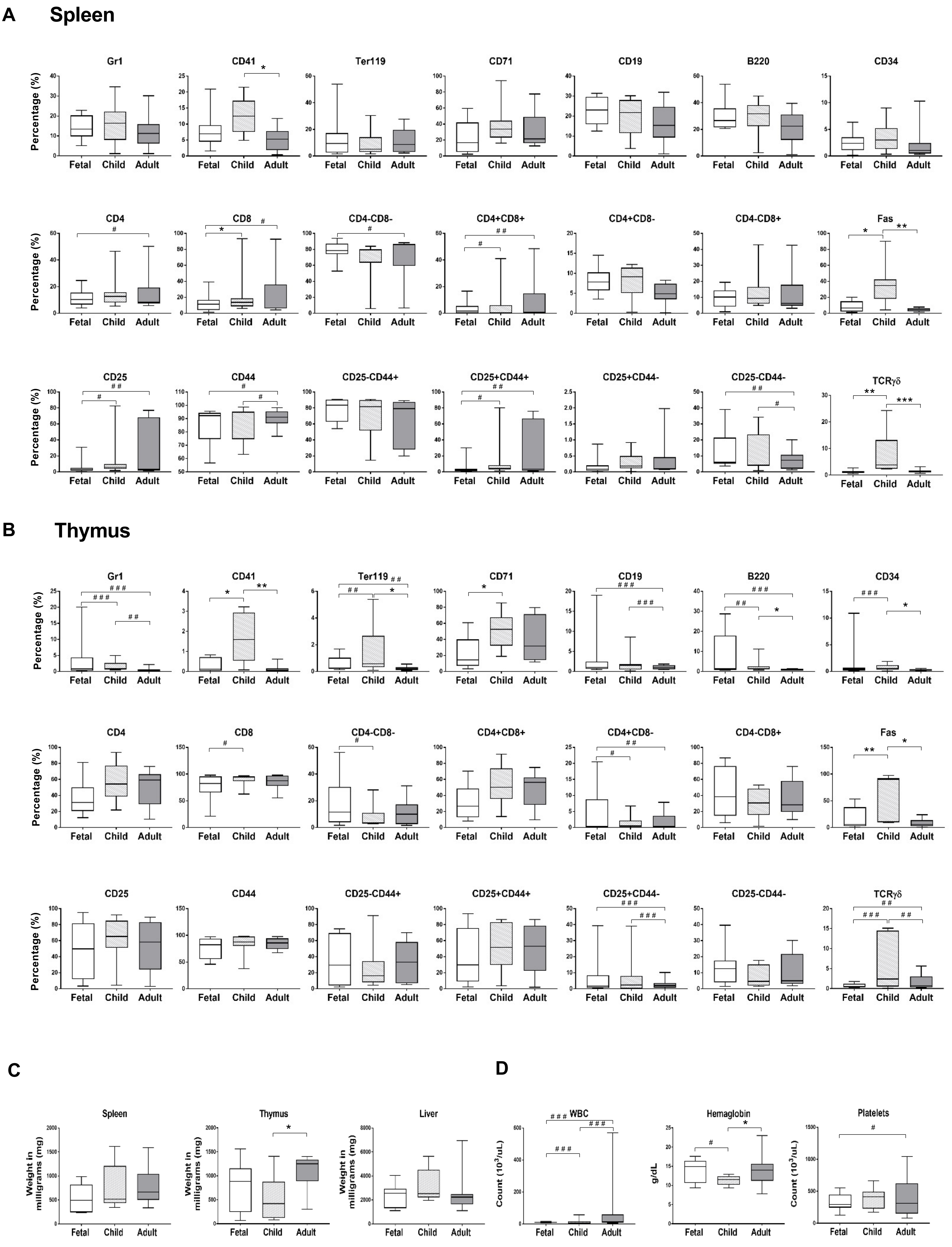
A
Spleen
Gr1
CD41
Ter119
CD71
CD19
B220
CD34
CD4
CD8
CD4-CD8-
CD4+CD8+
CD4+CD8-
CD4-CD8+
Fas
CD25
CD44
CD25-CD44+
CD25+CD44+
CD25+CD44-
CD25-CD44-
TCRγδ
Percentage (%)
Fetal
Child
Adult
B
Thymus
Gr1
CD41
Ter119
CD71
CD19
B220
CD34
CD4
CD8
CD4-CD8-
CD4+CD8+
CD4+CD8-
CD4-CD8+
Fas
CD25
CD44
CD25-CD44+
CD25+CD44+
CD25+CD44-
CD25-CD44-
TCRγδ
Percentage (%)
Fetal
Child
Adult
C
Spleen
Thymus
Liver
Weight in milligrams (mg)
D
WBC
Hemaglobin
Platelets
Count (10^3/uL)
g/dL
